# Multivariate Genome-wide and integrated transcriptome and epigenome-wide analyses of the Well-being spectrum

**DOI:** 10.1101/115915

**Authors:** B.M.L. Baselmans, R. Jansen, H.F. Ip, J. van Dongen, A. Abdellaoui, M. P. van de Weijer, Y. Bao, M. Smart, M. Kumari, G. Willemsen, J.J. Hottenga, BIOS consortium, Social Science Genetic Association Consortium, E.J.C. de Geus, D.I. Boomsma, M.G. Nivard, M. Bartels

## Abstract

Phenotypes related to well-being (life satisfaction, positive affect, neuroticism, and depressive symptoms), are genetically highly correlated (| *r*_*g*_ | > .75). Multivariate genome-wide analyses (*N*_*obs*_ = 958,149) of these traits, collectively referred to as the well-being spectrum, reveals 63 significant independent signals, of which 29 were not previously identified. Transcriptome and epigenome analyses implicate variation in gene expression at 8 additional loci and CpG methylation at 6 additional loci in the etiology of well-being. We leverage an anatomically comprehensive survey of gene expression in the brain to annotate our findings, showing that SNPs within genes excessively expressed in the cortex and part of the hippocampal formation are enriched in their effect on well-being.

---

Well-being plays an important role in psychology and medicine, as well as in economics^1,2^. Well-being owes its interdisciplinary prominence to its associations with physical and mental health, and its role as a desired socio-economic outcome and index of economic development^3^. Most existing research on the genetics of well-being is characterized by a focus on individual phenotypes, despite the strong correlations between related traits. The high genetic correlations (| *r*_*g*_ | > .75)^4^ between life satisfaction, positive affect, neuroticism, and depressive symptoms suggests common underlying biology, or a partially shared genetic etiology. Acknowledging this overlap, we performed a multivariate genome-wide meta-analysis (multivariate GWAMA) (*N*_*obs*_ = 958,149) of these four phenotypes to increase the power to identify associated genetic variants (**Supplementary Table 1**).

Our analyses leveraged published univariate GWAMA of life satisfaction^4,5^ (*N*_*obs*_ = 80,852), positive affect^4,5^ (*N*_*obs*_ = 189,028), neuroticism^4–6^ (*N*_*obs*_ = 238,315), and depressive symptoms^4,5,7,8^ (*N*_*obs*_ = 449,954). For the purpose of the multivariate GWAMA, we reversed the estimated SNP effects on neuroticism and depressive symptoms to ensure a positive correlation with life satisfaction and positive affect. The dependence between effect sizes (error correlation) induced by sample overlap was estimated from the genome-wide summary statistics obtained from the univariate GWAMA analyses using LD score regression (see online methods). Knowledge of the error correlation between univariate meta-analyses allowed dependent samples to be meta-analyzed, providing a gain in power while guarding against inflated type I error rates (see online methods).

We recognize that the measures included in the well-being spectrum are not necessarily interchangeable. Therefore, we performed two types of multivariate GWAMA; 1) N-weighted multivariate GWAMA, which assumes a single underlying construct (see online methods); 2) model averaging GWAMA, where we relaxed the assumption of a unitary effect of the SNP on all traits. For the latter analyses, we performed eight model-based GWAMA’s, capturing various combinations of SNP effects (see online methods). To account for within and between model variability, we took a weighted average of the effect size and standard error for each SNP across the eight models. For each of the eight models we computed an AICc weight to ensure that all models were weighted properly^9^.

In our N-weighted multivariate GWAMA, we identified 58 independent signals at 51 genomic loci (**Fig. 1A, Supplementary Table 2**). Our model averaging GWAMA detected five additional signals, bringing the total independent signals associated with the well-being spectrum to 63 (**Fig 1B-E, Supplementary Table 3-7**). Of these 63 signals, 29 were not identified in univariate analyses. In contrast, univariate analyses identified 53 unique signals, 18 were not significant in either the N-weighted multivariate GWAMA or the model averaging GWAMA (**Supplementary Fig. 1A-D**). Heterogeneity in terms of genome-wide significant signal may reflect imperfect genetic correlations, or reflect that the dichotomy between significant and not significant is arbitrary (e.g. 16 out of the 18 signals detected only in univariate analyses had a p-value below 10e-5 in our multivariate analyses). For both multivariate strategies we observed, in comparison to univariate GWAMA, higher median test statistics (N-weighted λ_GC_ = 1.533, average model-based λ_GC_ = 1.489, univariate GWAMA λ_GC_ = 1.407), but no increase in LD score intercept (N-weighted =1.003, average model-based = 0.934). The low LD score intercept confirmed that the inflation in test statistics was due to an increase in signal, rather than population stratification or inaccurate accounting for sample overlap (see online methods, **Supplementary Table 8, Supplementary Fig. 2-3**).

**Fig. 1.**
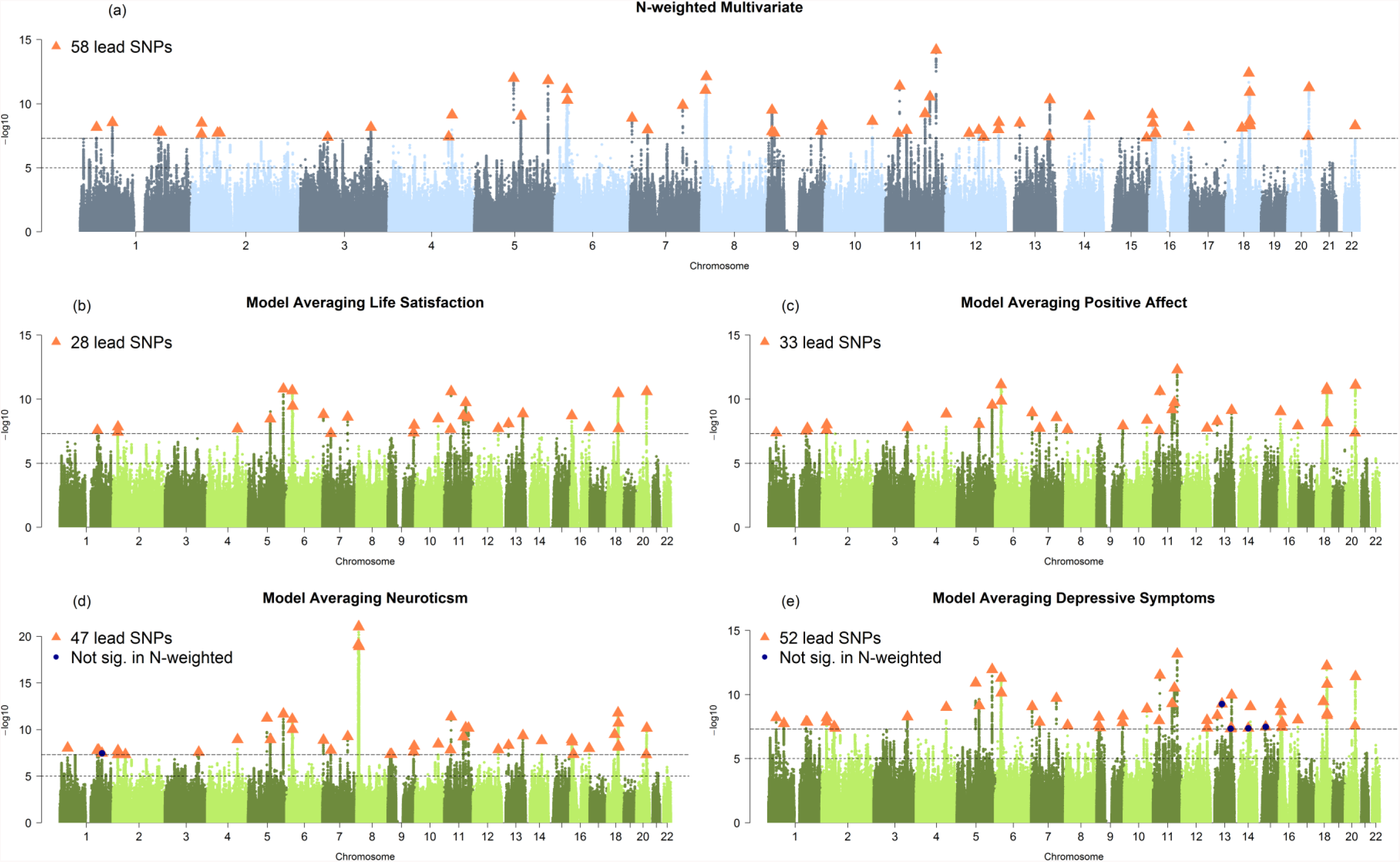
Manhattan plots of N-weighted and model averaging GWAMA. **(a)** N-weighted GWAMA and model averaging GWAMA of **(b)** life satisfaction, **(c)** positive affect, **(d)** neuroticism, **(e)** depressive symptoms. All plots in all panels are based on the same set of SNPs. The x-axis represents the chromosomal position, and the y-axis represents the significance on a −log10 scale. Each approximately independent genome-wide significant association (“lead SNP”) is marked by Δ.

To confirm the gain in power of our multivariate GWAMA, we performed polygenic risk score prediction (PRS) in two independent samples (combined sample size > 16,000; online methods)^5,10^. We predicted the phenotypes in the well-being spectrum (life satisfaction, positive affect, neuroticism, and depressive symptoms). The PRS based on the N-weighted GWAMA improved prediction with an average increase in R^2^ of 0.39 (1.51-fold change), where the increase in R^2^ ranged from 0.15 to 0.67 (i.e., a 1.13 to 4.60-fold change in R^2^). PRS based on model-average GWAMA revealed an average increase in R^2^ of 0.36 (i.e., a 1.48-fold change), which ranged from −0.01 to 0.67 (i.e., 0.99 to 4.60-fold change in R^2^, **Supplementary Fig. 4, Supplementary Table 9**).

Both flavors of multivariate GWAMA aggregated the effect of a single SNP across multiple traits, informed by prior knowledge of the genetic correlation between these traits. In a similar fashion, we proceeded to aggregate the effect across multiple SNPs based on prior knowledge that some of these SNPs collectively influence the expression level of a gene transcript or the methylation level at a CpG site. Aggregation of SNP effects, which have a common effect on gene expression or CpG methylation lowers the multiple testing burden, strengthens the signal, and identifies specific gene transcripts or CpG methylation sites. In this procedure, known as summary-based transcriptome-wide and methylome-wide association analyses (TWAS and MWAS)^11,12^, we imputed the effect of changes in gene expression or CpG site methylation on the well-being spectrum. Information on the relation between SNP and CpG methylation or gene expression were obtained from the summarized results of cis-expression (e)QTL and cis-methylation (m)QTL studies in whole blood^13,14^. Since there is evidence that cis-regulation of gene expression is highly consistent across tissues (genetic correlation brain-blood = .66)^15^, we used eQTL results obtained based on gene expression measure in blood as a proxy for expression in brain tissue. Given the equal performance of both multivariate GWAMAs and to avoid multiple testing, we used the results of the N-weighted GWAMA. We uncovered 34 transcript-trait associations, of which 8 are not identified in the GWAMA and are located more than 250kb from a significant GWAMA locus. We found 115 CpG methylation-trait associations mapping to 6 distinct loci (> 250kb from GWAMA loci) at a Bonferroni corrected significance level. When considering TWAS and MWAS loci significant at a Benjamini-Hockberg adjusted (FDR) p-value < 0.05^16^, we uncovered 121 transcript-trait associations, of which 105 are distinct (>250kb) and 859 CpG methylation sites associated with the well-being spectrum, mapping to 218 distinct novel loci more than 250kb from a significant GWAMA locus (**Supplementary Table 10-11**).

We performed further biological annotation using stratified LD score regression. We used 220 genomic annotations (33 brain and 187 non-brain annotations), which reflect four histone marks (H3K4me1, H3K4me3, H3K27ac, or H3K9ac) in 54 tissues. Our analyses revealed significant enrichment in 45 regions of the genome characterized by 30 histone marks in 10 brain tissues. Among these brain tissues are: mid frontal and inferior temporal lobe, fetal brain, cingulate and angular gyrus, germinal matrix (a highly cellular and highly vascularized region in the brain from which cells migrate out during brain development), hippocampus anterior caudate, substantia nigra, and the neurosphere (**Supplementary Table 12, Supplementary Fig. 5**). We found enrichment of SNP effects in a single endocrine tissue, the thymus. Note that other endocrine tissues, which are more frequently implicated in depression (e.g., the thyroid and pituitary gland) were not available for testing. We also found enrichment in 14 non-brain tissues (**Supplementary Fig. 5**). Further analyses using LD score regression, where the LD scores are based on evolutionary markers^17^, revealed that SNPs of very recent origin (i.e. a low allelic age) explained substantially more variation than ancient SNPs (**Supplementary Table 13 and Supplementary Fig. 6**). These findings indicate the presence of negative selection shaping variation in well-being through recent evolution (see online methods).

In order to more accurately pinpoint brain regions where genes relevant to the well-being spectrum are differentially expressed, we computed stratified LD scores based on differential gene expression in an anatomically comprehensive set of 210 brain regions, based on 3707 measurement in 6 human brains^18^. For each brain region, genes were selected that showed higher expression compared to all other regions, inclusion is determined based on a t-statistic of the difference test. For each region we selected the top 10% most strongly expressed genes. The LD scores were significantly enriched at FDR < 0.05 at multiple gyri in the cortex (e.g. fusiform gyrus, parahippocampal gyrus and precentral gyrus) as well as the precuneus, planum polare, temporal pole and the superior -and paracentral lobule (**Supplementary Table 14, Supplementary Fig. 7-13**). Differential gene expression appeared driven mainly by transcriptional differences between gross anatomical regions in the brain (cortex, sub-cortical structures, brainstem, and cerebellum). To reveal regions related to the well-being spectrum within these regions, we computed differential brain expression only *within* the cortex, sub-cortical structures, brainstem and cerebellum and identified enrichment of genes specifically expressed in the subiculum (Z = 3.60, p < 0.001; **Fig. 2, Supplementary Table 15**). The subiculum is considered part of the hippocampal formation and plays a key role in hippocampal-cortical interaction^19^ in the inhibition of the Hypothalamic-Pituitary-Adrenal-axis and the human response to stress^20^. As a negative control, we considered the enrichment of genes differentially expressed in all brain regions using height GWAMA summary statistics: no region was significantly enriched, both when considering either global or local differential expression (all p > 0.05). To test whether the signal observed in the subiculum was specific to the well-being spectrum, we repeated LD score regression analyses of genes differentially expressed in the subiculum for educational attainment^21^ and schizophrenia GWAMA summary statistics^22^. We found no enrichment of effect on educational attainment (Z = .11, p = .38) but did find an enrichment of effect on schizophrenia (Z = 3.22, p < 0.0007) for genes differentially expressed in the subiculum. All results of the differential gene expression analysis were mapped to the MNI coordinates at which the tissue samples were obtained, allowing for future integration of our findings and other neuroimaging modalities (**Supplementary Table 16**).

**Fig. 2.**
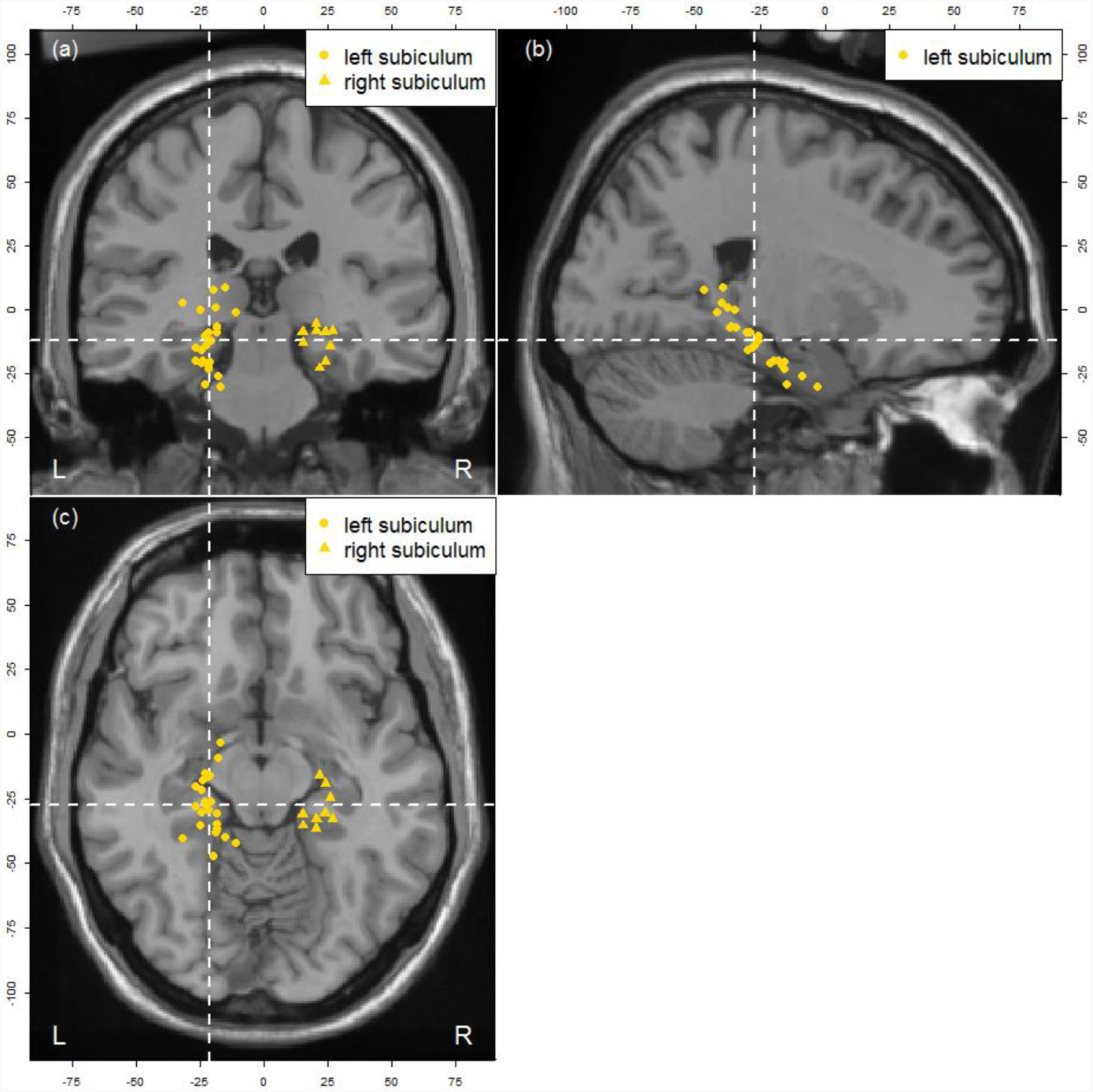
**Local differential gene expression between subcortical structures** identifies enrichment of genes specifically expressed in the subiculum (Z = 3.60, p < 0.001), in their effect on well-being. (a) coronial view (b) sagittal view (c) axial view. The sample location for brain tissues which were used to measured gene expression by Hawrylycz et al. (2012) is available and were projected to a standard MNI template brain (“Colin27”). The figure is centered on the averaged MNI coordinates of brains samples which are part of the annotation “left Subiculum” (x = 77, y = 90 and z = 60).

Gene expression LD scores driven biological annotations are based on empirical regional gene expression differences across the brain. The results therefore, by necessity, describes a mix of inter and intracellular processes that cannot be attributed to specific brain functions at the level of a cell or gene. To annotate individual genes identified in G/T/M-WAS loci, we performed a look-up off all genes in significant (187 genes) and suggestive loci (1512 genes) in the HGRI-EBI catalog^23^ of published genome-wide association studies. We found abundant evidence for possible pleiotropic effects (e.g. 56 genes shared with schizophrenia, 9 genes shared with educational attainment), but also genes that have not previously been associated with any trait (**Supplementary Table 17-19**). To shed light on cellular processes, we cross referenced the KEGG database^24^ and our genome-wide signals to previously suggested neurotransmission processes across the synaptic cleft (**Supplementary Table 20**). Three of the genome-wide significant loci contain genes involved in glutamatergic neurotransmission (*GNAO1*, *BRIK3*, *GRM5*) while another 8 genes, at suggestive associated loci (p < 10e-5 for GWAMA loci, FDR < 0.05 for T/M-WAS loci), are also involved in glutamatergic neurotransmission processes. Among these suggestive loci are *SLC1A7* and *SLC38A*, genes involved in the reuptake of glutamate from the synaptic cleft into the presynaptic terminals and glial cells. Several other genes are involved in post synaptic response to glutamate. We further found 2 genes at genome-wide significantly loci (*DDR2* and *GNAO1*) and another 10 genes at suggestive loci involved in dopaminergic neuro-transmission. Suggestive genes in the dopaminergic synapse pathway include *DRD2* and *DRD5* receptors. GNAO1 is further involved with GABAergic and serotonergic neurotransmission, as are another 11 and 9 genes respectively in suggestive loci. Among the suggestive serotonin related genes are the *HTR1E*, *HTR1D HTR3A* and *HTR3B* receptors, but not *5-HTT* or *HTR2A*, which have frequently been implicated in the etiology of depression and other psychiatric traits^25,26^. Among the suggestive signals in the GABAnergic synapse is the *GABBR1* gene which encodes for a gamma-aminobutric acid auto receptor involved in the inhibition of neurotransmitter release. When considering these genes in the context of synaptic neurotransmission, it is important to realize that many, if not all, of these genes serve functions in other pathways and biological mechanisms as well, and their primary effect on the well-being spectrum not necessarily arises from their effect on neurotransmission.

Another locus of particular interest was found within the major histocompatibility complex. Recent work has identified 3 individual signals related to schizophrenia in the MHC region, one of which is linked to complement 4 (C4A) gene expression and synapse elimination during puberty^27^. The genome-wide significant signal for the well-being spectrum in the MHC region is not in strong LD with lead eQTL’s for C4A gene expression. Rather a second independent signal tagged by rs13194504 is associated with both schizophrenia and well-being. TWAS and MWAS results for the MHC region implicate the expression of ZKSCAN4 and methylation of cg08798685 in the etiology of well-being (**Supplementary Fig. 14**).

In summary, while previous univariate analyses of phenotypes in the well-being spectrum were moderately successful, we gained power by use of multivariate analysis. Our work shows definitive progress in the genetics of well-being. Model averaging GWAMA identified additional loci, associated with some but not all traits in the well-being spectrum, and provided flexibility in terms of model specification. Model averaging can in fact incorporate *any* multivariate GWAMA or GWAS model for which the per SNP model fit can be expressed in terms of an AICc fit statistic. The averaging procedure is done per locus, allowing for heterogeneity across phenotype and loci. Additionally, we found that TWAS and MWAS can yet further increase the pool of loci related to variation in complex traits, like well-being. We accompany our multivariate analyses with genomic driven neuroimaging. By leveraging the genome-wide results, LD score regression, and an atlas of brain gene expression we were able to pinpoint brain regions where region specific gene expression exists for genes enriched in their effect on well-being, and we report evidence for enrichment of genes differentially expressed in several cortical regions, as well as in the subiculum. Our analyses are limited in the following ways: we consider regional brain expression, though the cause of regional differences in gene expression across the brain can reflect a host of underlying regional differences. Gene expression is known to vary systematically between cell-types within the brain^28^ (e.g neurons, microglia, astrocytes) and developmental phases^29^ (prenatally, childhood, adulthood and old age), and likely even between sub-types of a single cell type. Differences in gene expression across or within cell types may induce differences between regions as cell type composition may differ between regions. This limitation applies especially to the global enrichment of genes expressed in many cortical regions when compared to all other brain structures. However, the aforementioned limitations will be addressed in the future, as several efforts are underway to categorize gene expression across the human brain at increased (single cell) resolution. Single cell sequencing (e.g. drop-seq based anatomically comprehensive survey of the brain), based on donors deceased at different ages, could disentangle cell type specific from region specific differential gene expression as well as age specific gene expression^30^. Our study maps the results of a GWAMA of well-being to brain regions based on a coordinate system shared with multiple other neuroscientific measurements, facilitating future integration between the genetic study of - and neuroscientific research on the well-being spectrum.

## Online Methods

### N-weighted multivariate GWAMA

We obtained summary statistics from previous published studies^4–8^, where multiple cohorts contributed to the univariate GWAMAs of life satisfaction, positive affect, neuroticism and depressive symptoms http://www.thessgac.org/. To quantify the dependence between the univariate GWAMAs, we estimated the cross trait LD score intercept (CTI)^31,32^.

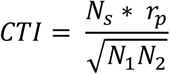

Where N_s_ equals the sample overlap, N_1_ the sample size for trait one and N_2_ the sample size for trait two, *r*_*p*_ equals the phenotypic correlation between trait one and two. The CTI is approximately equal to the covariance between the test statistics obtained in a GWAMA of trait 1 and trait 2. We assume the estimated CTI is equal to the true CTI, though note the uncertainty in the estimated CTI is generally low. Given the estimated covariance between effect sizes we can meta-analyse the four dependent GWAMAs and obtain a multivariate test statistic per SNP:

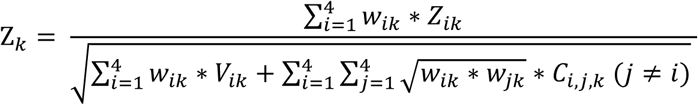

Where w_ik_ is the square root of the sample size for SNP k in the GWAMA of trait i, Z_ik_ is the test statistics of SNP k in the GWAMA of trait i; V_ik_ is the variance of the test statistic for SNP k in the GWAMA of trait i (i.e 1 given that Z is a standardized test statistic) and C_i,j,k_ is the covariance between test statistics for SNP k between GWAMA of trait i and trait j (where C equals CTI obtained from cross trait LD score regression between trait i and trait j). The multivariate test statistic *Z*_*k*_, is a standardized sum of tests statistics all of which follow a normal distribution under their respective null distributions, the statistic *Z*_*k*_ follows a standard normal distribution under the null hypothesis of no effect.

### Model averaging GWAMA

Consider the following model:

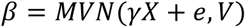

Where *β* (1xn) is a multivariate normal vector of effect sizes obtained from the regression of n standardized phenotypes on a standardized genotype (SNP). The matrix V (nxn) is the variance-covariance matrix of effect sizes, matrix X a design matrix (pxn), and *γ* the corresponding vector of parameters (1xp). The indexed p denotes the number of variables included in the means model of the response vector *β*.

In this context, a regular GWAMA restricts the design matrix X to a unit vector (i.e. we model a single genetic effect, which is assumed identical across cohorts, and any observed variation is attributed to sample fluctuation). Generally, matrix V is diagonal, and contains the squared standard errors of elements in *β*. A regular GWAMA is the most restricted model one can consider. However, when considering multivariate GWAMA (i.e. the elements in β reflect SNP effects on separate yet correlated phenotypes) this model might be too restrictive even when traits have a substantial genetic correlation, not all genetic effects need to be shared between traits or be identical in magnitude. The least restrictive model is to consider the SNP effects in *β* independent (i.e. run univariate GWAMA of the correlated phenotypes). In between the most restrictive and least restrictive model, a manifold of models can be specified, equating
the effects in y across combinations of traits, while allowing it to differ between other combinations of traits. These models can be specified by ways of the design matrix X.

One could consider a manifold (z) of models (m), each with a different design matrix X.

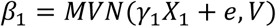

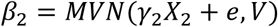

…

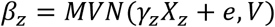

When considering i correlated phenotypes, a simple expansion of X is to allow for 2 vectors (p = 2), a unit vector and a second vector which is coded dichotomously (0,1) where the coding varies over each of the m models. Other codings, based on analysis of the genetic correlation between traits (i.e. PCA or Cholesky decomposition), can be applied to summary statistics and included in the average. Practically, this allows for the existence of 2 distinct genetic effect. This procedure results in .5 * *k*^2^ models. The 1df model with a unit vector for X and .5 * *k*^2^ – 1 2-df models with a unit vector and a second vector which codes for all possible combinations of pairs of k traits. However, simply considering m models for all SNPs across the genome results in a prohibitive increase of the already substantial multiple testing burden. Given m possible models, each of which predict a different vector *γ*, and uncertainty for the predicted elements in *γ*, a possible way forward is to average the model predictions. The models are weighted by the relative proportion of evidence for each model. Specifically, the weights can be based on the AICc^33^ information criteria. The AICc for model m equals:

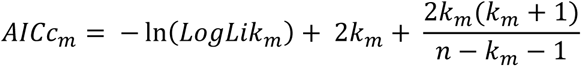

For each AICc we compute the delta (Δ_m_) to the best (i.e lowest) AICc value, and from these we compute the model weights (g) for the k models as:

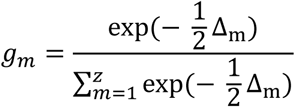

We predict the vector β using each of the models

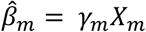

One can aggregate the prediction over all models as:

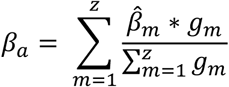

And we aggregate the uncertainty within and between models to obtain *var* (*β*_*a*_):

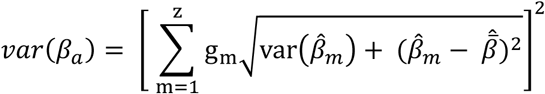

The resulting vector *β*_*a*_ contains the model averaged effect sizes for the effect of a particular SNP on the phenotypes subjected to multivariate analysis. Note how the variance estimate contains a variance component which reflects within model variability 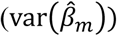 which equals the square of the standard error, and a variance component between model variability 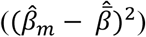 in estimate, which ensures no overfitting occurs.

Our procedure boosts power if the SNP effect is concordant between traits, while retaining strongly discordant SNP effects if the model favors these. Model averaging offers several avenues for extension. One can constrain the SNP effects across multiple SNPs based on biological knowledge of the relation between the SNPs and gene expression, or CpG methylation (analog to TWAS). Alternative it might be beneficial to average the AICc weights across regions of the genome. Model averaging can in principle accommodate *any model* for which the AICc information criterion can be expressed. These models should result in a vector of SNPS effects (*β*) and an asymptotic variance for the SNP effects. In the current application, models per SNP are estimated in R using the “metafor” package and models are averaged using the “AICcmodavg” package^34,35^.

### Polygenic Risk Prediction

To confirm the gain in power of our multivariate GWAMA results, we perform polygenic risk score prediction (PRS) in two independent samples; 1) the Netherlands Twin Register (NTR)^1010,36^ and 2) Understanding Society (UKHLS)^5^. We predict the phenotypes in the well-being spectrum (life satisfaction, positive affect, neuroticism, and depressive symptoms). In NTR, LS and PA data are available in 9,143 and 6,836 genotyped participants, respectively. LS is measured longitudinally using the Satisfaction with Life Scale consisting of five items (e.g., “My life is going more or less as I wanted”) with responses given on a seven-point scale, resulting in a minimum score of five and a maximum score of 35^37^. PA is also measured longitudinally using four questions that were adapted from the Subjective Happiness Scale^38^ (e.g., “On the whole, I am a happy person”) with responses on a seven-point scale, resulting in a minimum score of four and a maximum score of 28. Neuroticism data are available for 8,527 genotyped participants. The Big Five personality traits (including neuroticism) were measured by using the NEO-FFI^39^, a sixty-item personality questionnaire consisting of five subscales: neuroticism, extraversion, openness, agreeableness and conscientiousness. The responses were given on a five-point scale (0-4). Subscale scores are constructed for each time point by taking the sum across the twelve subscale-specific items (after recoding opposite-stated items), and are set to missing if ten or more items of the total scale are unanswered. When subjects have fewer than ten missing items, missing items are scored at two (which is neutral given the 0-4 scale). Depressive symptoms are obtained from the DSM-oriented Depression subscale of the age-appropriate survey from the ASEBA taxonomy^40^ and are available in 7,898 participants. To measure depressive symptoms, fourteen questions are used (e.g., “Enjoys little”) and responses were given on a three-point scale ranging from zero (“not true”) to two (“very true”). The DSM-oriented subscale is constructed for each time point by taking the sum across the fourteen subscale-specific items and is set to missing if more than twenty percent of the total survey items were unanswered. When less than twenty percent of items are missing for a participant, the missing items are replaced by the participant’s mean score.

In UKHLS data is available in 9,944 participants. LS was measured longitudinally (waves 1-6). Participants were asked how satisfied they were “with life overall” with responses given on a seven-point scale, resulting in a minimum score of one and a maximum score of seven. PA is also measured longitudinally (waves 1 and 4 only) using The Warwick-Edinburgh Mental well-being scale (WEMWBS). SWEMWBS is a shortened version of WEMWBS. This 7-item short version (see Tennant et al., 2007) is scored on a 5-point Likert scale, from “none of the time” to “all of the time”, and summed to give a total score, ranging from 7 to 35 Neuroticism data are available for 8,198 genotyped participants from wave 3. The Big Five personality traits (including neuroticism) were measured using The Big Five Inventory (BFI), a 44-item personality questionnaire consisting of five subscales: neuroticism, extraversion, openness, agreeableness and conscientiousness. The responses were given on a seven-point scale (1-7). The neuroticism score combines three items on the neuroticism subdomain. Component scores were calculated as the average item response if no more than one of the three input responses was missing. Depressive symptoms (DS) were measured longitudinally (waves 1-6) and obtained from The General Health Questionnaire (GHQ) available in 9,203 participants. The 12 question GHQ was used containing questions relating to concentration, loss of sleep and general happiness. The 12 questions are scored on a four-point scale (1-4). Valid answers to the 12 questions of the GHQ-12 were converted to a single scale by recoding 1 and 2 values on individual variables to 0, and 3 and 4 values to 1, and then summing, giving a scale running from 0 (the least distressed) to 12 (the most distressed).

The weights used for the polygenic scores are based on our two flavors of multivariate GWAMAs as well as the four univariate GWAMAs. Scores are based on the intersection of SNPs available in any of these GWAMAs. In the NTR, SNPs were imputed to a common reference SNPs with MAF < 0.005, Hardy-Weinberg Equilibrium (HWE) with *p* < 10^−12^ and call rate < 0.95 were removed. Individuals are excluded from the analyses if the genotyping call rate is < 0.90, the inbreeding coefficient as computed in PLINK48 (F) was < −0.075 or > 0.075), if the Affymetrix Contrast QC metric is < .40, if the Mendelian error rate is > 5 standard deviations (SDs) from the mean, or if the gender and Identity-by-State (IBS) status does not agree with known relationship status and genotypic assessment. In UKHLS, SNPs were imputed to a common reference (1000 Genomes project March 2012 version 3). SNPs with MAF < 0.01, HWE *p* < 10^−4^ and call rate < 0.98 were removed, individuals with A B and C were removed In NTR 1,224,793 SNPs passed QC and were used to construct polygenic scores and in UKHLS 955,441 SNPs passed QC and were used to construct polygenic scores. The phenotypes were regressed on sex and age as well as principal components which were included to correct for ancestry and the polygenic scores. Results can be found in **Supplemental Table 9**.

### Summary-Based transcriptome wide (TWAS) and methylome wide (MWAS) association studies

We used the tool DIST^41^ to impute the HapMap reference based results for the N-weighted GWAMA to the 1000Genomes Phase1 reference. We aggregate SNP effects informed by their common effect on expression level of gene or CpG methylation, as was proposed by Gusev et al.^11^ We used the BIOS eQTL resource as eQTL reference set to build imputation models to predict gene expression using multiple eQTL SNPs^13^. Models are built per gene (gene models) by identifying independent eQTL SNPs based on stepwise conditional regression.^13^ The z-score for each eQTL SNP is used in TWAS as a weight (q).The eQTLs used are available at http://genenetwork.nl/biosqtlbrowser/. Based on the gene models, N-weighted GWAMA summary statistics and LD based on the GONL reference^42^, TWAS is performed. That is, for each gene-prediction-model containing eQTLs S_1_- S_N_ with weights q=q_1_,q_2_,…,q_n_, the corresponding GWAMA z-scores z=z_1_,z_2_,…,z_n_ and LD is an n-by-n correlation matrix for eQTLs S_1_- S_N_, were used to
construct a test statistic:

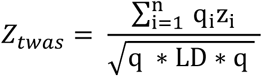

MWAS was performed following the same procedure to build imputation models to predict CpG site methylation of the DNA strand using multiple mQTL SNPs. The methylation site specific weights were obtained from the BIOS mQTL study^14^.

### Stratified LD score regression

To determine whether specific genomic regions are enriched for genetic effects on the well-being spectrum phenotypes, we used LD Score regression^31,32^. We are specifically interested in regions of the genome which are histone modified in a specific tissue. For example, regions of the genome which are histone modified in the prefrontal cortex, can be transcribed more frequently in prefrontal tissue. The enrichment of these genomic regions in their effect on well-being suggest the involvement of processes in the prefrontal cortex in the etiology of wellbeing.

LD Score regression is based on the relationship between the observed chi-square of a SNP and the degree of LD between a SNP and its neighbor. SNPs in strong LD are more likely to tag causal effects on complex traits and therefore have a higher expected chi-square. The procedure can be extended to *stratified* LD score regression where multiple LD scores are created, each of which captures the LD for a SNP with other SNPs of a specific category of interest, for example SNPs in a histone modified region of the genome.

We follow the exact procedure described by Finucane et al.^43^ We estimated stratified LD Score regression for the “baseline” model, which contains 53 categories. The model consists of a category containing all SNPs, 24 categories corresponding to main annotations of interest, 24 categories corresponding to 500-bp windows around the main annotations, and categories corresponding to 100-bp windows around ChIP-seq peaks (i.e. regions that are Sensitive to DNase1 or associated with histones bearing the modification marks H3K4me1, H3K4me3, H3K27ac or H3K9ac). In addition to the analysis of the baseline model, we performed analyses using cell type-specific annotations for the four histone marks, which correspond to specific chemical modifications to the histone protein, which in turn package and orders the DNA molecule. Epigenetic modifications of histones, specifically histones bearing the marks H3K4me1, H3K4me3, H3K27ac or H3K9ac, all of which are associated with increased transcription of DNA into RNA. Each cell type-specific annotation corresponds to a histone mark in a specific cell obtained from distinct human tissue, for example H3K27ac in Fetal Brain cells, generating 220 combinations of histone modification by tissue. When generating estimates of enrichment for the 220 Histone mark by tissue annotations, we control for overlap with the functional categories in the full baseline model, but not for overlap with the 219 other cell type specific annotations. Then for our well-being phenotype, we ran LD Score regression on each of the 220 models (one for each histone by tissue combination) and ranked the histone by tissue annotations by P-value derived from the Z-values of the coefficient. Results are displayed in **Supplementary Table 12**.

### Stratified LD score regression to detect negative selection

Using LD score regression, where the LD scores are based on evolutionary markers, we test whether negative selection is present in shaping variation in the well-being spectrum through recent evolution. We use the same procedure as extensively described by Gazal et al.^17^ and report the proportion of heritability for each LD-related annotation of the baseline model subdivided in 5 quantiles ranging from 20% common SNPs with youngest allelic age to 20% of common SNPs with oldest allelic age. These proportions are computed based on a joint fit of the baseline-LD model, but measure the heritability explained by each quantile of each annotation while including the effect of other annotations. Results are displayed in **Supplementary Table 13**. The effect of nucleotide diversity, recombination rate, background selection, CpG-content predicted allelic age, and levels of LD in Africa all are consistent with those found in Gazal et al., in which Gazal et al show by ways of forward simulation to be consistent with the presence of negative selection^17^.

### Stratified LD score regression of local gene expression across the human brain

We downloaded the normalized and QC’ed gene expression measured in an anatomically comprehensive set of brain regions from http://www.brain-map.org/. The data contains 3707 measurements across 6 adult human brains. The procedures used to measure gene expression and standardize these measures across the brains are described in Hawrylycz et al^18^. Based on these data we compute differential gene expression for 48154 probes which map to 20724 unique genes (probes which did not map to genes were omitted). We considered differential gene expression across 210 regions for which at least 3 measurements are available. As Hawrylycz et al.^18^ found little evidence for lateral difference in gene expression, regions in the left and right hemisphere are collapsed into a single region. For each gene in each region a t-test is performed, testing the difference in standardized expression between the region in question and all other brain regions. Top 10% of probes ranked in terms of t-statistic per region a were retained. The unique genes mapped to this set of probes was extracted (mapping ∼2900-3500 genes to each region). The correlation between t-statistics for the 48154 probes revealed fairly strong differential expression between the cortex, brainstem and cerebellum and clustering of differential expression within these regions.

A partitioned LD score with respect to the genomic regions spanned by these genes (using gencode v19 as a reference), and a 100 kilobase window around each gene, is computed. The heritability of well-being is partitioned across the 54 baseline annotations developed by Finucane et al^43^ and each of the 210 brain regions, the regions are considered separately. The substantial differences in gene expression between gross anatomical brain regions (cerebellum, cortex, sub-cortical regions and brainstem) dominate the results (**Supplemental Fig. 7-13, Supplementary Table 14**). We therefore proceed to compute differential gene expression within the cerebellum, cortex, sub cortical regions and brainstem. In this analysis we omit the fibre bundles as these are anatomically distinct from both the cortex and the sub cortical regions, yet not measured densely enough to warrant the computation of differential expression within the fibre bundle tissues. The procedure to compute differentially expressed genes is identical to the procedure used to compute differential expression across the whole brain, but considers the gross anatomical regions separately. New LD scores are computed based on the local differential gene expression analysis and the resulting enrichment is visualized in **Figure 2 (Supplementary Table 15)**. All analyses were repeated using height as a negative control phenotype. The genomic regions spanning genes differentially expressed in these 210 brain regions are not significantly enriched with SNP effects on height.

### GWAS Catalog lookup

We search the NHGI GWAS catalog^23^ to determine which of our genome-wide significant GWAMA, MWAS, and TWAS signals have been previously reported (database searched on 03-04-2017). We apply two strategies; 1) for our GWAMA analyses, we identify genes that are within a 250kb distance from a genome wide significant SNPs For our TWAS and MWAS analyses the focus is specifically on markers that were not significant in GWAMA analyses. Therefore, we rule out any genes that we identify in TWAS or near MWAS hits that are within 250kb distance from a GWAMA lead SNP. Gene transcripts and genes near CpG sites which conform to the criteria outlined above are looked up in the GWAS catalog. Results can be found in **Supplementary Table 17-19**.

### Intersection between genome wide signals and synaptic pathways

Using the KEGG database^24^, all genes in the four synaptic pathways; (1) serotonergic synapse (2) dopaminergic synapse (3) glutamatergic synapse and (4) GABAergic synapse, we extracted. The four resulting gene sets where intersected with genes residing in the 77 genome wide significant loci. Similary, we looked at the intersection between the four synaptic pathways and suggestive loci (SNPs associated with our multivariate GWAMA at p < 5 * 10^−5^ and FDR < 0.05 for T/M-WAS loci, **Supplementary Table 20**).

## Acknowledgements

We would like to thank all participants in the cohort studies. This work was supported by the Netherlands Organization for Scientific Research (NWO: MagW/ZonMW grants 904-61-090, 985-10-002,904-61-193,480-04-004, 400-05-717, NWO-bilateral agreement 463-06-001, NWO-VENI 451-04-034, Addiction-31160008, Middelgroot-911-09-032, Spinozapremie 56-464-14192), Biobanking and Biomolecular Resources Research Infrastructure (BBMRI –NL, 184.021.007), the VU University’s Institute for Health and Care Research (EMGO^+^) and Neuroscience Campus Amsterdam (NCA), the European Science Council (ERC Advanced, 230374), the Avera Institute for Human Genetics, Sioux Falls, South Dakota (USA) and the National Institutes of Health (NIH, R01D0042157-01A). Part of the genotyping was funded by the Genetic Association Information Network (GAIN) of the Foundation for the US National Institutes of Health (NIMH, MH081802) and by the Grand Opportunity grants 1RC2MH089951-01 and 1RC2 MH089995-01 from the NIMH. Part of the analyses were carried out on the Genetic Cluster Computer (http://www.geneticcluster.org), which is financially supported by the Netherlands Scientific Organization (NWO 480-05-003), the Dutch Brain Foundation, and the department of Psychology and Education of the VU University Amsterdam. M.Bartels is/was financially supported by a senior fellowship of the (EMGO^+^) Institute for Health and Care and a VU University Research Chair position. MGN is supported by Royal Netherlands Academy of Science Professor Award (PAH/6635) awarded to DIB and supported by Biobanking and Biomolecular Resources Research Infrastructure (BBMRI–NL, 184.021.007.).

Understanding Society is an initiative funded by the Economic and Social Research Council (ES/H029745/1) and various Government Departments, with scientific leadership by the Institute for Social and Economic Research, University of Essex, and survey delivery by NatCen Social Research and Kantar Public. The research data are distributed by the UK Data Service. The genome-wide scan data were analysed and deposited by the Wellcome Trust Sanger Institute. Information on how to access the data can be found on the Understanding Society website https://www.understandingsociety.ac.uk/. Genotype-phenotype data access for UKHLS is available by application to Metadac: www.metadac.ac.uk.

## Individual Author Contribution

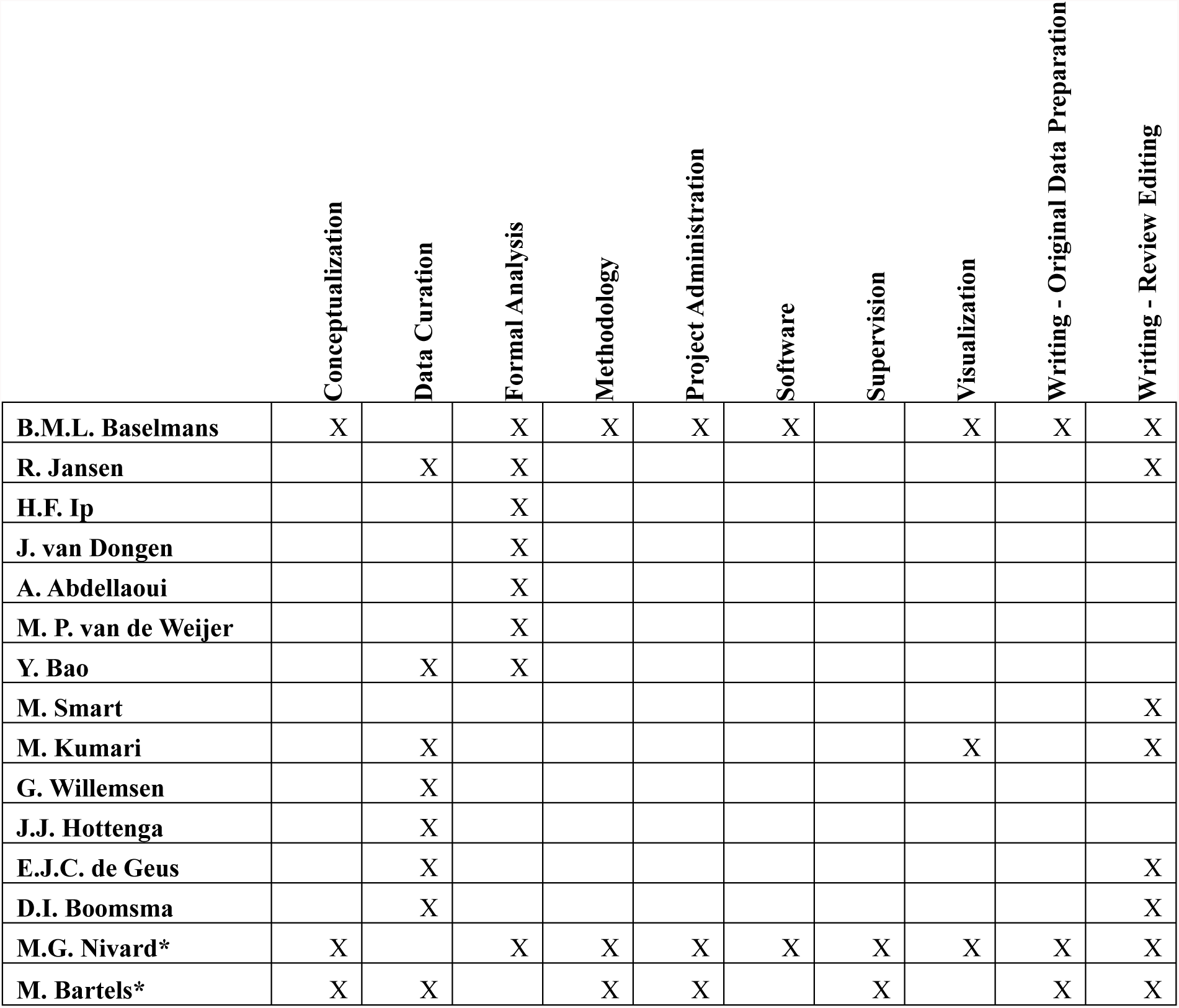

## Figure legends

**Supplementary Figure 1.** Manhattan plots of univariate GWAMA. **(a)** life satisfaction, **(b)** positive affect, **(c)** neuroticism, **(d)** depressive symptoms. The *x*-axis represents the chromosomal position, and the *y*-axis represents the significance on a −log_10_ scale. Each approximately independent genome-wide significant association (“lead SNP”) is marked by Δ.

**Supplementary Figure 2.** Quantile-quantile plots for the N-weighted and 4 average GWAMAs. **(a)** is the N-weighted GWAMA. **(b)** is model-average life satisfaction, **(c)** model average positive affect, **(d)** model average neuroticism **(e)** model average depressive symptoms

**Supplementary Figure 3.** Quantile-quantile plots for the four univariate GWAMAs. **(a)** life satisfaction, **(b)** average positive affect, **(c)** neuroticism **(d)** depressive symptoms

**Supplementary Figure 4.** The result of polygenic risk prediction based on univariate discovery GWAMA, N-weighted discovery GWAMA or model averaging discovery GWAMA. The unit on the Y-axis is the R-squared in percentage, obtained from a regression of the trait on the PRS, age, sex and 10 principle components. **(a)** displays the combined N-weighted polygenic score results. **(b)** displays the polygenic prediction results from the Netherlands Twin Register and **(c)** displays the polygenic results from Understanding Society. LS is life satisfaction. PA is positive affect. NEU is neuroticism. DEP is depressive symptoms.

**Supplementary Figure 5.** 220 Cell specific histone modified region enrichment. The bar plot is reflecting the FDR adjusted p-value for tissue specific histone modified regions of the genome, as estimated using partitioned LD-score regression. Blue bars represent brain regions, black bars represent non-brain regions.

**Supplementary Figure 6.** Proportion of heritability explained by quintiles of each LD-related annotation. The red line indicates the proportion of heritability when there is no enrichment (20% of SNPs explain 20% of heritability.

**Supplementary Figure 7.** Global differential gene expression in all 210 cortical structures identifies enrichment of genes specifically expressed in the **(1)** cingulate gyrus, inferior, **(2)** cingulate gyrus, superior, **(3)** angular gyrus, superior, **(4)** fusiform gyrus, bank of cos, **(5)** fusiform gyrus, bank of the its and **(6)** fusiform gyrus, lateral bank of the gyrus. For all structures, the Z -and p-values for the differential gene expression as well as the MNI coordinates are provided. To determine the MNI coordinates, we averaged the mni xyz coordinates of the left brain structures separately.

**Supplementary Figure 8.** Global differential gene expression in all 210 cortical structures identifies enrichment of genes specifically expressed in the **(1)** Heschl’s gyrus, **(2)** inferior temporal gyrus, bank of the its, **(3)** inferior temporal gyrus, lateral bank of the gyrus, **(4)** inferior temporal gyrus, bank of mts, **(5)** long insular gyri and **(6)** Lingual gyrus. For all structures, the Z -and p-values for the differential gene expression as well as the MNI coordinates are provided. To determine the MNI coordinates, we averaged the mni xyz coordinates of the left brain structures separately.

**Supplementary Figure 9.** Global differential gene expression in all 210 cortical structures identifies enrichment of genes specifically expressed in the **(1)** middle frontal gyrus, inferior, **(2)** middle frontal gyrus, superior, **(3)** medial orbital gyrus, **(4)** middle temporal gyrus, inferior, **(5)** middle temporal gyrus, superior and **(6)** Occipito-tempral gyrus, superior. For all structures, the Z -and p-values for the differential gene expression as well as the MNI coordinates are provided. To determine the MNI coordinates, we averaged the mni xyz coordinates of the left brain structures separately.

**Supplementary Figure 10.** Global differential gene expression in all 210 cortical structures identifies enrichment of genes specifically expressed in the **(1)** Paracentral lobule, anterior part, **(2)** Precuneus, inferior, **(3)** Precuneus, superior, **(4)** Parahippocampal gyrus, bank of cos, **(5)** Parahippocampal gyrus, lateral bank of the gyrus and **(6)** Planum polare. For all structures, the Z -and p-values for the differential gene expression as well as the MNI coordinates are provided. To determine the MNI coordinates, we averaged the mni xyz coordinates of the left brain structures separately. To determine the MNI coordinates, we averaged the mni xyz coordinates of the left brain structures separately.

**Supplementary Figure 11.** Global differential gene expression in all 210 cortical structures identifies enrichment of genes specifically expressed in the **(1)** Postcentral gyrus, central sulcus, **(2)** Postcentral gyrus, posterior of the central sulcus, **(3)** Precentral gyrus, central sulcus, **(4)** Precentral sulcus, inferior lateral aspect of the sulcus, **(5)** Precentral gyrus, bank of the precentral sulcus and **(6)** Superior frontal gyrus. For all structures, the Z -and p-values for the differential gene expression as well as the MNI coordinates are provided. To determine the MNI coordinates, we averaged the mni xyz coordinates of the left brain structures separately.

**Supplementary Figure 12.** Global differential gene expression in all 210 cortical structures identifies enrichment of genes specifically expressed in the **(1)** Superior frontal gyrus, medial bank, **(2)** Supramarginal gyrus, inferior, **(3)** Supramarginal gyrus, superior, **(4)** Superior occipital gyrus, **(5)** Superior parietal lobule and **(6)** Superior temporal gyrus. For all structures, the Z -and p-values for the differential gene expression as well as the MNI coordinates are provided. To determine the MNI coordinates, we averaged the mni xyz coordinates of the left brain structures separately.

**Supplementary Figure 13.** Global differential gene expression in all 210 cortical structures identifies enrichment of genes specifically expressed in the **(1)** Transverse gyri, **(2)** Temporal pole, inferior and **(3)** Temporal pole, superior. For all structures, the Z -and p-values for the differential gene expression as well as the MNI coordinates are provided. To determine the MNI coordinates, we averaged the mni xyz coordinates of the left brain structures separately.

**Supplementary Figure 14.** Local association in the MHC region. (a) provides a local Manhattan plot for the MHC region with interposed on top the LD with a strong eQTL for the C4 gene linked to neuronal pruning in adolescence and schizophrenia by Sekar et al^27^. (b) is a scatter plot for the −log_10_(p) against the R2 with the C4 eQTL. (c) provides a local Manhattan plot for the MHC region with interposed on top the LD with SNP rs13194504, the strongest MHC signal found for schizophrenia. (d) is a scatter plot of the – log_10_(p) against the R2 with rs13194504. Round symbols represent SNPs, square symbols represent gene transcripts and triangle symbols represent CpG sites.

## MEMBERS OF SOCIAL SCIENCE GENETIC ASSOCIATION CONSORTIUM

The following people, who are not listed as coauthors on this manuscript, contributed to the original GWAMA on Subjective Well-being, Depressive Symptoms, and Neuroticism, on which the present paper is based. The views presented in the present paper may not reflect the opinions of the individuals listed below.

We thank: Aysu Okbay, Jan-Emmanuel De Neve, Patrick Turley, Mark Alan Fontana, S Fleur W Meddens, Richard Karlsson Linnér, Cornelius A Rietveld, Jaime Derringer, Jacob Gratten, James J Lee, Jimmy Z Liu, Ronald de Vlaming, Tarunveer S Ahluwalia, Jadwiga Buchwald, Alana Cavadino, Alexis C Frazier-Wood, Nicholas A Furlotte, Victoria Garfield, Marie Henrike Geisel, Juan R Gonzalez, Saskia Haitjema, Robert Karlsson, Sander W van der Laan, Karl-Heinz Ladwig, Jari Lahti, Sven J van der Lee, Penelope A Lind, Tian Liu, Lindsay Matteson, Evelin Mihailov, Michael B Miller, Camelia C Minica, Ilja M Nolte, Dennis Mook-Kanamori, Peter J van der Most, Christopher Oldmeadow, Yong Qian, Olli Raitakari, Rajesh Rawal, Anu Realo, Rico Rueedi, Börge Schmidt, Albert V Smith, Evie Stergiakouli, Toshiko Tanaka, Kent Taylor, Gudmar Thorleifsson, Juho Wedenoja, Juergen Wellmann, Harm-Jan Westra, Sara M Willems, Wei Zhao, LifeLines Cohort Study, Najaf Amin, Andrew Bakshi, Sven Bergmann, Gyda Bjornsdottir, Patricia A Boyle, Samantha Cherney, Simon R Cox, Gail Davies, Oliver S P Davis, Jun Ding, Nese Direk, Peter Eibich, Rebecca T Emeny, Ghazaleh Fatemifar, Jessica D Faul, Luigi Ferrucci, Andreas J Forstner, Christian Gieger, Richa Gupta, Tamara B Harris, Juliette M Harris, Elizabeth G Holliday, Jouke-Jan Hottenga, Philip L De Jager, Marika A Kaakinen, Eero Kajantie, Ville Karhunen, Ivana Kolcic, Meena Kumari, Lenore J Launer, Lude Franke, Ruifang Li-Gao, David C Liewald, Marisa Koini, Anu Loukola, Pedro Marques-Vidal, Grant W Montgomery, Miriam A Mosing, Lavinia Paternoster, Alison Pattie, Katja E Petrovic, Laura Pulkki-Råback, Lydia Quaye, Katri Räikkönen, Igor Rudan, Rodney J Scott, Jennifer A Smith, Angelina R Sutin, Maciej Trzaskowski, Anna E Vinkhuyzen, Lei Yu, Delilah Zabaneh, John R Attia, David A Bennett, Klaus Berger, Lars Bertram, Dorret I Boomsma, Harold Snieder, Shun-Chiao Chang, Francesco Cucca, Ian J Deary, Cornelia M van Duijn, Johan G Eriksson, Ute Bültmann, Eco J C de Geus, Patrick J F Groenen, Vilmundur Gudnason, Torben Hansen, Catharine A Hartman, Claire M A Haworth, Caroline Hayward, Andrew C Heath, David A Hinds, Elina Hyppönen, William G Iacono, Marjo-Riitta Järvelin, Karl-Heinz Jöckel, Jaakko Kaprio, Sharon L R Kardia, Liisa Keltikangas-Järvinen, Peter Kraft, Laura D Kubzansky, Terho Lehtimäki, Patrik K E Magnusson, Nicholas G Martin, Matt McGue, Andres Metspalu, Melinda Mills, Renée de Mutsert, Albertine J Oldehinkel, Gerard Pasterkamp, Nancy L Pedersen, Robert Plomin, Ozren Polasek, Christine Power, Stephen S Rich, Frits R Rosendaal, Hester M den Ruijter, David Schlessinger, Helena Schmidt, Rauli Svento, Reinhold Schmidt, Behrooz Z Alizadeh, Thorkild I A Sørensen, Tim D Spector, John M Starr, Kari Stefansson, Andrew Steptoe, Antonio Terracciano, Unnur Thorsteinsdottir, A Roy Thurik, Nicholas J Timpson, Henning Tiemeier, André G Uitterlinden, Peter Vollenweider, Gert G Wagner, David R Weir, Jian Yang, Dalton C Conley, George Davey Smith, Albert Hofman, Magnus Johannesson, David I Laibson, Sarah E Medland, Michelle N Meyer, Joseph K Pickrell, Tõnu Esko, Robert F Krueger, Jonathan P Beauchamp, Philipp D Koellinger, Daniel J Benjamin & David Cesarini

## MEMBERS OF THE BIOS CONSORTIUM

The following people, who are not listed as coauthors on this manuscript, contributed to the collection and analysis of the expression and methylation data on which the present paper is based. The views presented in the present paper may not reflect the opinions of the individuals listed below.

### Management Team

Bastiaan T. Heijmans (chair), Peter A.C. ’t Hoen, Joyce van Meurs, Aaron Isaacs, Rick Jansen, Lude Franke

### Cohort collection

Dorret I. Boomsma, René Pool, Jenny van Dongen, Jouke J. Hottenga (Netherlands Twin Register); Marleen MJ van Greevenbroek, Coen D.A. Stehouwer, Carla J.H. van der Kallen, Casper G. Schalkwijk (Cohort study on Diabetes and Atherosclerosis Maastricht); Cisca Wijmenga, Lude Franke, Sasha Zhernakova, Ettje F. Tigchelaar (LifeLines Deep); P. Eline Slagboom, Marian Beekman, Joris Deelen, Diana van Heemst (Leiden Longevity Study); Jan H. Veldink, Leonard H. van den Berg (Prospective ALS Study Netherlands); Cornelia M. van Duijn, Bert A. Hofman, Aaron Isaacs, André G. Uitterlinden (Rotterdam Study).

### Data Generation

Joyce van Meurs (Chair), P. Mila Jhamai, Michael Verbiest, H. Eka D. Suchiman, Marijn Verkerk, Ruud van der Breggen, Jeroen van Rooij, Nico Lakenberg.

### Data management and computational infrastructure

Hailiang Mei (Chair), Maarten van Iterson, Michiel van Galen, Jan Bot, Dasha V. Zhernakova, Rick Jansen, Peter van ’t Hof, Patrick Deelen, Irene Nooren, Peter A.C. ’t Hoen, Bastiaan T. Heijmans, Matthijs Moed.

### Data Analysis Group

Lude Franke (Co-Chair), Martijn Vermaat, Dasha V. Zhernakova, René Luijk, Marc Jan Bonder, Maarten van Iterson, Patrick Deelen, Freerk van Dijk, Michiel van Galen, Wibowo Arindrarto, Szymon M. Kielbasa, Morris A. Swertz, Erik. W van Zwet, Rick Jansen, Peter-Bram ’t Hoen (Co-Chair), Bastiaan T. Heijmans (Co-Chair).

